# Task- and experience-dependent modulation of head-eye coordination during ball interception

**DOI:** 10.64898/2026.08.20.746110

**Authors:** Saya Ogino, Tomohiro Kizuka, Seiji Ono

## Abstract

Head-eye coordination during ball interception depends on both task demands and motor experience. The purpose of this study is to determine how these factors influence head-eye contributions to gaze control. Twenty-five female university students (novices with no ball sport experience, *n* = 13; experienced softball players, *n* = 12) performed two tasks: visually tracking an approaching ball (tracking task) and, in addition, moving the hand to the ball’s landing position (reaching task). Head, eye, and gaze velocities, cross-correlation coefficients between gaze and head velocity, and gaze-head lag time were analyzed using linear mixed models. The results showed that although gaze velocity remained unchanged regardless of tasks or groups, decomposing gaze into head and eye components revealed task-dependent contributions.

Compared with the tracking task, the reaching task showed significantly larger head velocity and smaller eye velocity, indicating complementary adjustments that were not revealed by gaze movements alone. The cross-correlation between head and gaze was significantly higher in the reaching task than the tracking task, indicating stronger temporal coupling under greater task demand. Furthermore, the experienced group showed greater task-dependent modulation of eye velocity than the novice group, demonstrating greater flexibility in adjusting the magnitude of head-eye movements to task demands. In addition, the experienced group showed a consistently near-zero gaze-head lag regardless of task, whereas the novice group showed a prolonged gaze-leads-head interval. These findings suggest that ball sport experience shapes two distinct aspects of head-eye coordination: task-dependent flexibility in movement magnitude, and stable, temporally synchronized gaze-head control.

## 1. Introduction

Ball interception tasks, such as catching and batting, are complex motor behaviors that require predicting the trajectory of an approaching ball while moving the body into an appropriate position. One-handed catching has been distinguished into two components: an orientation phase, in which the hand is directed toward the ball’s destination and requires spatial precision, and a flexion phase, in which the fingers close to secure the ball and requires temporal precision. Successful catching depends on accurately predicting the ball’s motion from information sampled relatively early in its flight [1]. Similarly, in prehension tasks more broadly, the transportation component, which directs the hand toward the target location, and the manipulation component, which grasps the object, have been shown to be executed in temporal coordination [2]. Because these reaching and grasping movements cannot be performed without accurately predicting where the ball will arrive, visual tracking of the approaching ball is essential.

Smooth pursuit eye movements play a functional role in responding to an approaching ball. Smooth pursuit accuracy has been shown to strongly predict interception accuracy and strategy in an interception task involving baseball players [3]. Predictive saccades toward the anticipated location of the ball have also been reported in ball sports such as cricket [4,5]. Furthermore, smooth pursuit eye movements have been shown to enhance prediction of visual motion [6], and visual uncertainty has been shown to systematically affect both eye and hand movements during interception [7]. However, most laboratory studies examining gaze behavior during interception tasks have been conducted with the head restrained [3,8], and head movements were not measured.

In head-free conditions, gaze in space is expressed as the sum of head rotation in space and eye rotation relative to the head [9,10]. When tracking a moving ball under head-free conditions, head rotation provides an angular acceleration stimulus to the semicircular canals, which in turn evokes a vestibulo-ocular reflex (VOR) that drives the eyes in the opposite direction. Although the VOR normally stabilizes gaze during head movement, when tracking a moving target it acts to divert gaze away from the target, requiring VOR suppression [9,11]. In gaze control involving such VOR suppression, the eye and head can each be controlled by independent predictive mechanisms, and it has been demonstrated experimentally that head and gaze can be controlled simultaneously at completely independent frequencies [12], suggesting that, given the relationship gaze = head + eye, the coordination of head and eye is not uniformly determined by a single control mechanism but can flexibly vary according to task demands. Moreover, brainstem pursuit pathways integrate visual, vestibular, and proprioceptive signals during combined eye-head tracking [13], and the coordination of eye, head, and hand movements has been examined during a continuous interception task [14]. Therefore, to comprehensively understand gaze behavior during ball interception, it is necessary to examine the coordinative relationship between the head and eye.

During ball interception, it has been shown that orientation signals from both the eye and head are required to integrate visual and proprioceptive information for reaching and grasping movements [15,16], and it is predicted that head involvement will be greater during reaching movements than during visual tracking alone. In sports involving interceptive actions, cross-correlation analysis has been used to quantify the temporal coupling among gaze, head, and arm movements [17]. Regarding the influence of ball sport experience on head-eye movement coordination, novice players have been shown to display greater peak head and eye velocities than experienced softball players, whereas experienced players show a gaze strategy that stabilizes gaze by predicting the ball trajectory [18]. In addition, the visuomotor delay in interceptive tasks has been shown to be shorter in experts than in novices [19], and skilled batters have been shown to initiate the directional change of head movement earlier and with less variability than novices [20]. However, these studies have examined either the speed of visuomotor processing or the timing of a single movement component relative to an external event, rather than the temporal coordination between head movement and the resulting gaze. Whether such head-gaze temporal coordination extends to tasks involving reaching toward or simulated catching of an approaching ball, as opposed to visual tracking alone, and how such coordination and its timing differ with ball sport experience, remain unclear. Quantifying the temporal coordination of head and gaze would allow a more precise examination of the characteristics of predictive gaze-head control in experienced players.

Based on the above, the present study aimed to determine head-eye movement coordination during tracking of an approaching ball from the perspectives of task demands and ball sport experience. The hypotheses were: (1) head velocity would be greater in the condition involving a reaching movement than in the condition involving visual tracking alone; (2) novice players would show greater head and eye velocities than experienced players; and (3) experienced players would show a more predictive temporal coordination pattern than novices, characterized by a smaller temporal lag between gaze and head.

## 2. Materials and Methods

### 2.1 Participants

Twenty-five right-handed female university students (mean age 20.0 ± 1.2 years) participated in this study. Participants were divided into two groups based on the presence and duration of ball sport experience. The experienced group comprised 12 female members of a university women’s softball team with at least 5 years of competitive experience in softball or baseball (mean age 20.7 ± 1.2 years; softball/baseball experience: 7.4 ± 2.3 years; total ball sport experience: 9.0 ± 2.7 years). The novice group comprised 13 participants with no history of formal ball sport instruction through club or extracurricular activities (mean age 19.4 ± 0.8 years; ball sport experience: 0 years). This study was conducted in accordance with the Declaration of Helsinki (2013 revision), and all experimental protocols were approved by the Research Ethics Committee of the Institute of Health and Sport Sciences, University of Tsukuba (approval number: 025-53; approved October 29, 2025). All participants received a full explanation of the study purpose and procedures prior to participation and provided written informed consent.

### 2.2 Experimental setup

A schematic diagram of the experimental setup is shown in Fig. 1. A tennis ball launcher (Tennis Tutor Plus, Sports Tutor, Fig. 1-①) projected tennis balls toward an acrylic board positioned in front of the launcher (Fig. 1-②). The ball outlet of the launcher was set at a height of 125 cm above the ground, and the distance to the participant (Fig. 1-③) was 9 m. The initial ball velocity was maintained at approximately 40 km/h (11.1 m/s) throughout the experiment. To measure head and eye movements during the experimental tasks, participants wore a wearable eye tracker (Pupil Invisible or Pupil Neon, Pupil Labs, Fig. 1-④), and calibration was performed prior to measurement. Head movements were recorded using the gyroscope sensor built into the eye tracker. For the Pupil Invisible, both head and eye movements were sampled at 200 Hz. For the Pupil Neon, head movements were sampled at 110 Hz and eye movements at 200 Hz. To synchronize the timing of ball launch with the eye tracker data, a photoelectric sensor placed at the ball outlet (Fig. 1-⑤) triggered an LED light (Fig. 1-⑥) at the moment of each launch. The LED activation was captured in the scene video (30 Hz) recorded from the participant’s perspective by the eye tracker. Synchronization was achieved by manually identifying the frame in which the LED activated in the scene video and aligning the corresponding scene video timestamp with the wearable eye tracker data timestamp [21].

**Fig. 1.**
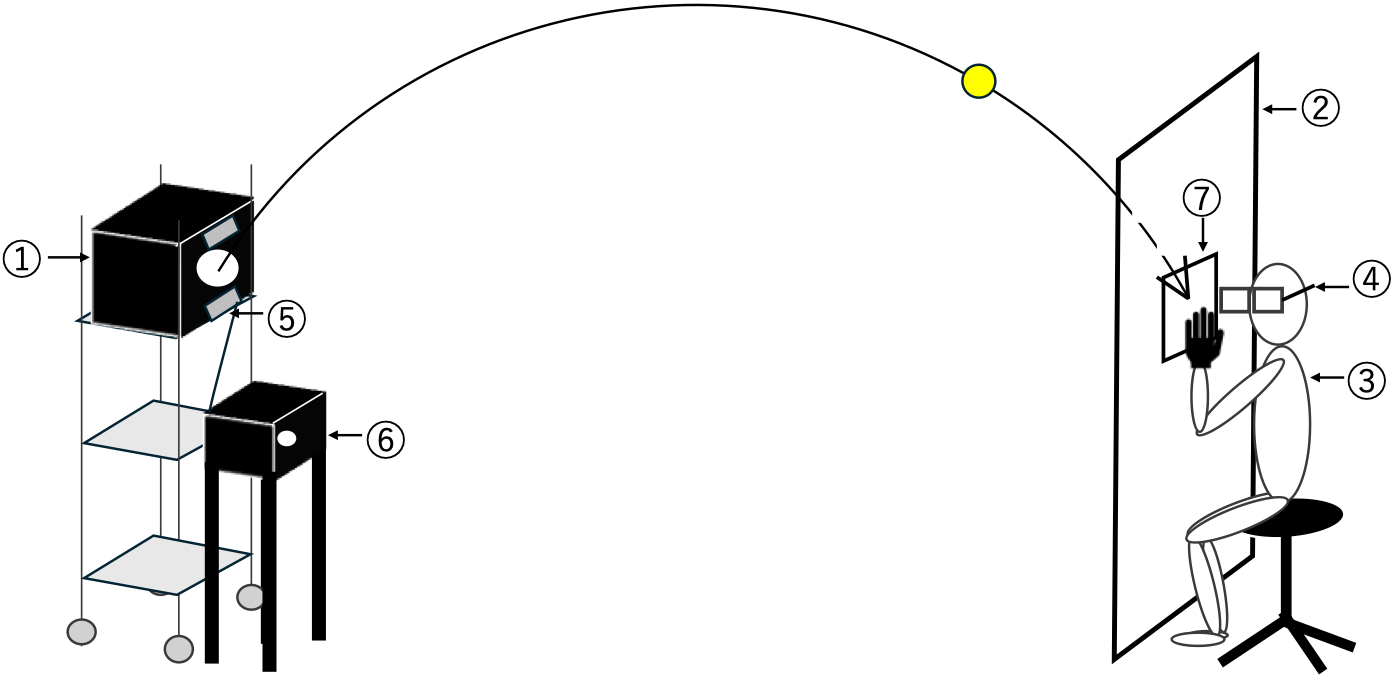
Schematic diagram of the experimental setup: ①Tennis ball launcher, ②Acrylic board, ③ Participant, ④Wearable eye tracker, ⑤Photoelectric sensor, ⑥LED light, ⑦Designated landing zone for incoming balls.

### 2.3 Experimental tasks

Two experimental tasks were used in this study: (1) a task requiring participants to visually track an approaching ball (tracking task), and (2) a task requiring participants to move their hand to the position where the ball arrived on the acrylic board (reaching task). During both tasks, participants sat on a chair adjusted to eye height 125 cm above the ground. Trials were repeated until 20 balls had landed within the designated zone on the acrylic board (Fig. 1-⑦; 30 × 30 cm).

In the tracking task, participants were instructed to visually track the ball continuously from launch until it hit the acrylic board (Fig. 1-②). The mean ball flight time was 973.2 ms. The acrylic board was positioned 20 cm in front of the participant, who tracked the ball through the transparent acrylic board.

In the reaching task, participants were instructed to move their left hand to the position where the ball would arrive on the acrylic board (without actual ball contact). The left (non-dominant) hand was used to simulate a catching motion with the glove hand, as in softball and baseball catching with the non-throwing hand. In each trial, participants rested their hand on the acrylic board in a set position and, upon ball launch, slid it horizontally along the surface of the board to place the ball’s arrival position. The mean ball flight time was 970.1 ms. Task order was counterbalanced across participants.

### 2.4 Data analysis

From the data recorded by the eye tracker, time-series data for head pitch angle and eye elevation angle in the vertical direction were extracted. Each trial was defined as the interval from ball launch (trigger onset) to the moment the ball contacted the acrylic board (trigger offset), and data were extracted accordingly. Head, eye, and gaze positions at the moment of ball launch were set as the baseline (0 deg) for normalization. A fourth-order Butterworth low-pass filter with a cutoff frequency of 50 Hz was applied to the eye position data prior to saccade detection [10]. Eye velocity (deg/s) was calculated by differentiating eye position (deg) and used to detect the first saccade occurring after ball launch. Saccades were defined as rapid eye movements in which eye velocity exceeded 50 deg/s and persisted for at least 20 ms. A 400 ms analysis window was then defined with reference to the offset of this initial saccade, beginning 20 ms after saccade offset and ending 420 ms after saccade offset. Trials in which the full 400 ms window could not be secured, or in which missing data exceeded 20% of the window, were excluded from analysis. To extract the smooth tracking (slow-phase) component within the analysis window, saccades were removed, and the corresponding data were replaced by linear interpolation. The interpolated eye position data were then filtered using a fourth-order Butterworth low-pass filter with a cutoff frequency of 30 Hz [18]. The same 30 Hz filter was applied to the head position data. Because the raw sampling timestamps varied across trials and were not necessarily aligned to a fixed grid, the filtered head and eye position data were resampled onto a common 200 Hz grid (5-ms intervals) using linear interpolation. Gaze position was computed as the angular sum of the resampled head and eye position data. Head, eye, and gaze velocities in the vertical direction were then obtained by differentiating each position data. For each trial, it was verified that no blink occurred at the time of ball launch and that the ball landed within the designated zone on the acrylic board.

To evaluate head-eye coordination during the tracking phase of each trial, the 400 ms window was selected to capture the smooth pursuit phase following the initial saccade, while minimizing the inclusion of subsequent predictive saccades that appeared in some trials beyond this interval (Fig. 2). Trials in which the full 400 ms window could not be secured were excluded from analysis. Mean velocity of gaze, head, and eye movements in the vertical direction was calculated over the analysis window. To examine the temporal relationship between gaze and head movements in each task, the cross-correlation coefficient (CC) and the lag time between gaze and head velocities were computed. CC was defined as the maximum positive value of the cross-correlation function between gaze and head velocities. Lag time was defined as the time shift at which the cross-correlation function reached its maximum positive value (CC).

**Fig. 2.**
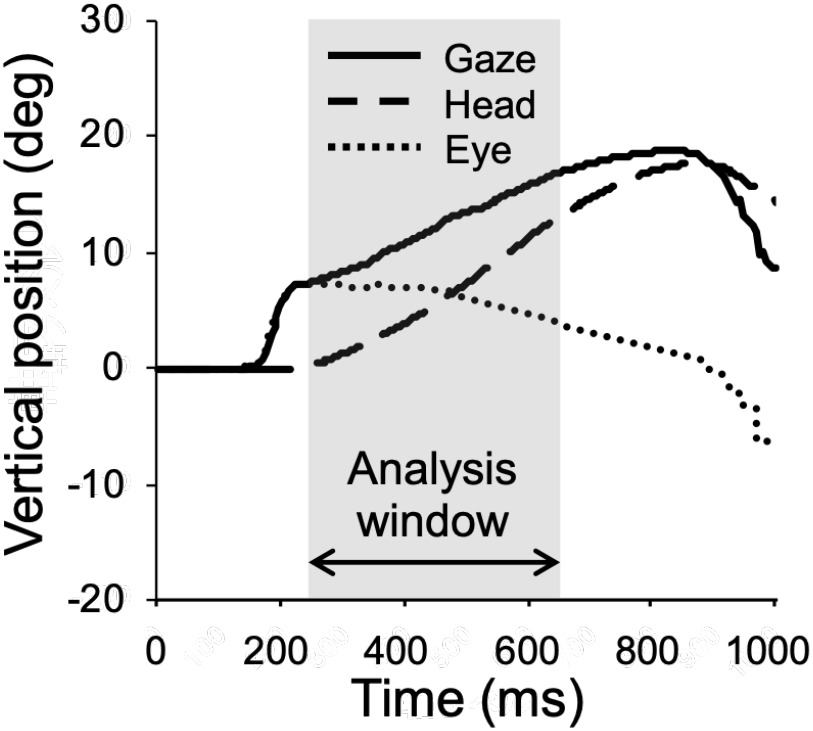
An illustrative example of gaze, head, and eye position from ball launch to contact with the acrylic board, showing the analysis window: t = 0 indicates the moment of ball launch. Mean ball flight time was 973.2 ms in the tracking task and 970.1 ms in the reaching task. Gaze, head, and eye positions at the moment of ball launch were normalized to baseline (0 deg). Solid, dashed, and dotted lines indicate gaze, head, and eye position, respectively. The shaded area indicates the 400 ms analysis window, defined as 20 to 420 ms after the offset of the initial saccade.

### 2.5 Statistical analysis

Linear mixed models (LMMs) were used to analyze each dependent variable, with trial-level data as the unit of analysis. Group, Task, and the Group × Task interaction were entered as fixed effects, and participant was included as a random intercept. When the interaction was significant, simple effects tests were conducted with Bonferroni correction. Effect sizes were calculated as partial eta-squared (η^2^p) from Type III F-statistics and degrees of freedom. All statistical analyses were performed using IBM SPSS Statistics version 30 (SPSS Inc., IL, USA). The significance level was set at p < .05 for all analyses. All data are presented as estimated marginal means ± standard error.

## 3. Results

### 3.1 Task-dependent changes in head and eye velocity

Representative time-series data of head and eye position are shown in Fig. 3. In both groups, head position showed greater movements in the reaching task than in the tracking task, whereas eye position moved more in the tracking task than in the reaching task.

**Fig. 3.**
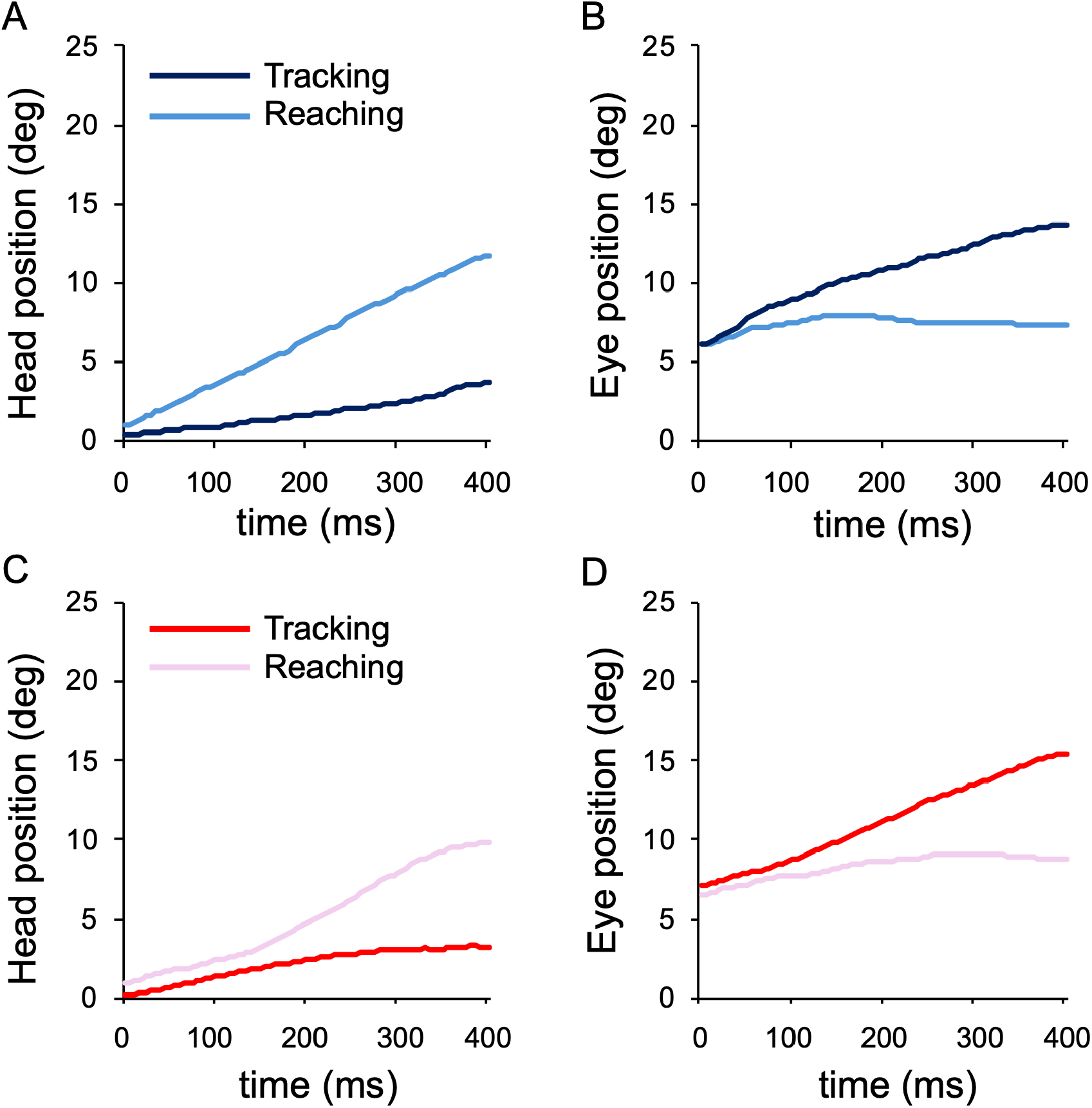
Representative examples of head and eye position time series: (A) Trial-averaged head position of a representative novice participant. (B) Trial-averaged eye position of a representative novice participant. (C) Trial-averaged head position of a representative experienced participant. (D) Trial-averaged eye position of a representative experienced participant. Each panel shows trial-averaged waveforms for the tracking task (dark blue or dark red) and reaching task (light blue or pink) over the 400 ms analysis window. t = 0 indicates the onset of the analysis window.

LMM results for head velocity showed a significant main effect of task (*F* (1, 608.07) = 201.03, *p* < .001, η^2^p = .25). Head velocity was significantly larger in the reaching task (19.36 ± 1.63 deg/s) than in the tracking task (10.71 ± 1.63 deg/s) (Fig. 4A). The main effect of group (*F* (1, 27.03) = 4.07, *p* = .054, η^2^p = .13) and the Task × Group interaction (*F* (1, 608.07) = 1.79, *p* = .182, η^2^p = .003) were not significant. LMM results for eye velocity showed a significant main effect of task (*F* (1, 604.59) = 136.87, *p* < .001, η^2^p = .185). Eye velocity was significantly larger in the tracking task (13.24 ± 1.57 deg/s) than in the reaching task (5.47 ± 1.56 deg/s) (Fig. 4B). The main effect of group was not significant (*F* (1, 23.28) = 2.14, *p* = .157, η^2^p = .084).

**Fig. 4.**
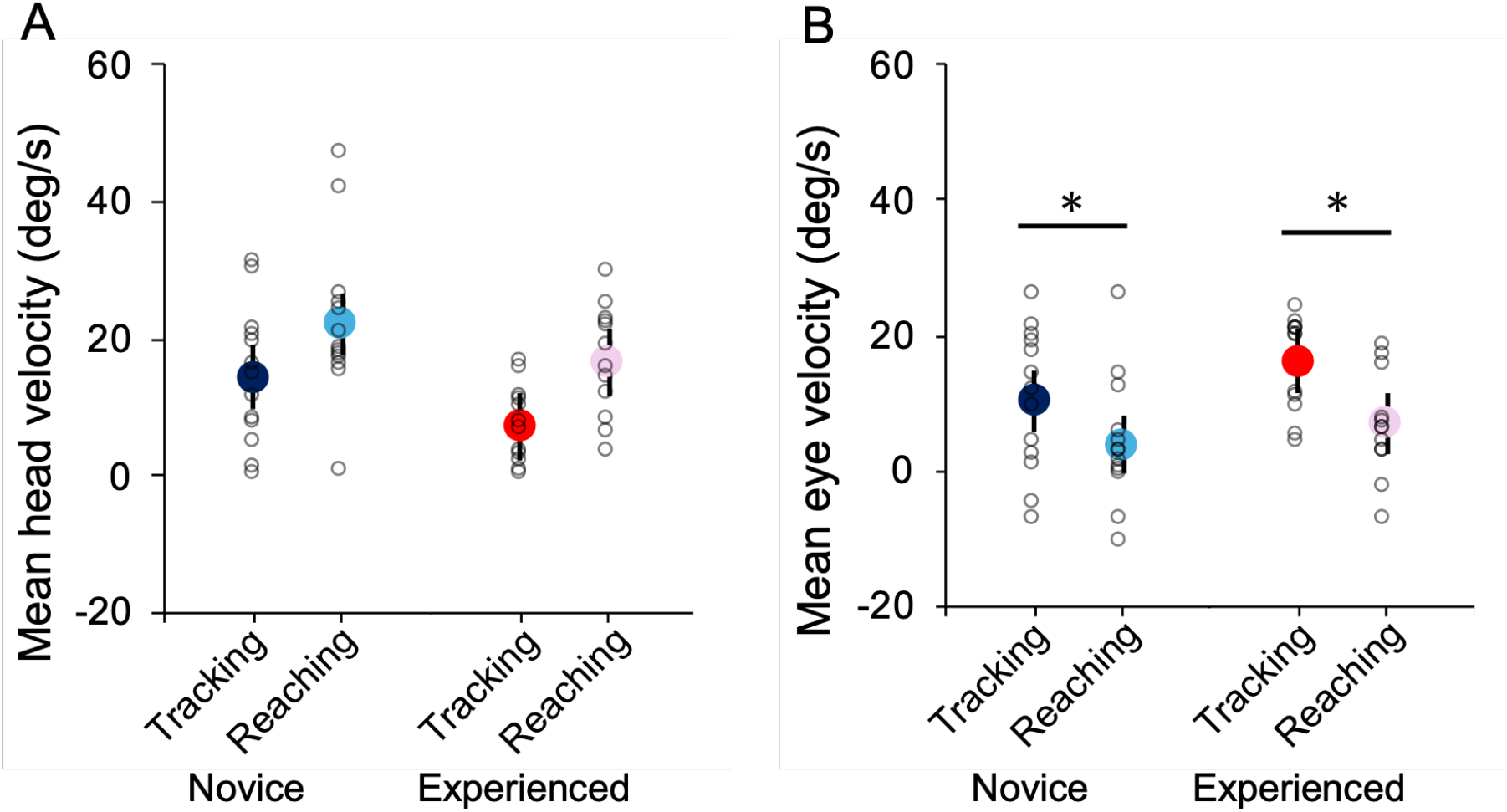
Head and eye velocity across tasks and groups: (A) Comparison of mean head velocity between tasks in each group. (B) Comparison of mean eye velocity between tasks in each group. Filled circles represent estimated marginal means; error bars indicate standard error. Open circles represent individual participant means, each averaged across that participant’s valid trials. Dark blue: novice group, tracking task; light blue: novice group, reaching task; dark red: experienced group, tracking task; pink: experienced group, reaching task. * *p* < .05.

The Task × Group interaction was significant (*F* (1, 604.59) = 4.25, *p* = .040, η^2^p = .007), indicating that the task-dependent change in eye velocity differed between groups, with the experienced group showing a greater reduction in eye velocity from the tracking to the reaching task than the novice group. Simple effects tests for task confirmed that eye velocity was significantly larger in the tracking task than in the reaching task in both the novice group (tracking: 10.31 ± 2.15 deg/s, reaching: 3.92 ± 2.16 deg/s; *p* < .001) and the experienced group (tracking: 16.16 ± 2.27 deg/s, reaching: 7.03 ± 2.26 deg/s; *p* < .001), although the reduction was larger in the experienced group. Simple effects tests for group showed no significant between-group differences in either the tracking task (*p* = .074) or the reaching task (*p* = .329).

### 3.2 Gaze velocity is unaffected by task demands or ball sport experience

Representative time-series data of gaze position are shown in Fig. 5. In contrast to the opposite directional changes in head and eye velocity between tasks, gaze velocity showed a similar pattern across tasks in both groups.

LMM results for gaze velocity showed no significant main effect of task (*F* (1, 605.13) = 3.47, *p* = .063, η^2^p = .006), no significant main effect of group (*F* (1, 23.15) = 1.29, *p* = .268, η^2^p = .053), and no significant Task × Group interaction (*F* (1, 605.13) = 1.24, *p* = .267, η^2^p = .002) (Fig. 6). No significant differences were observed between the tracking task (24.03 ± 0.92 deg/s) and the reaching task (24.92 ± 0.91 deg/s), or between the novice group (25.48 ± 1.22 deg/s) and the experienced group (23.47 ± 1.28 deg/s).

**Fig. 5.**
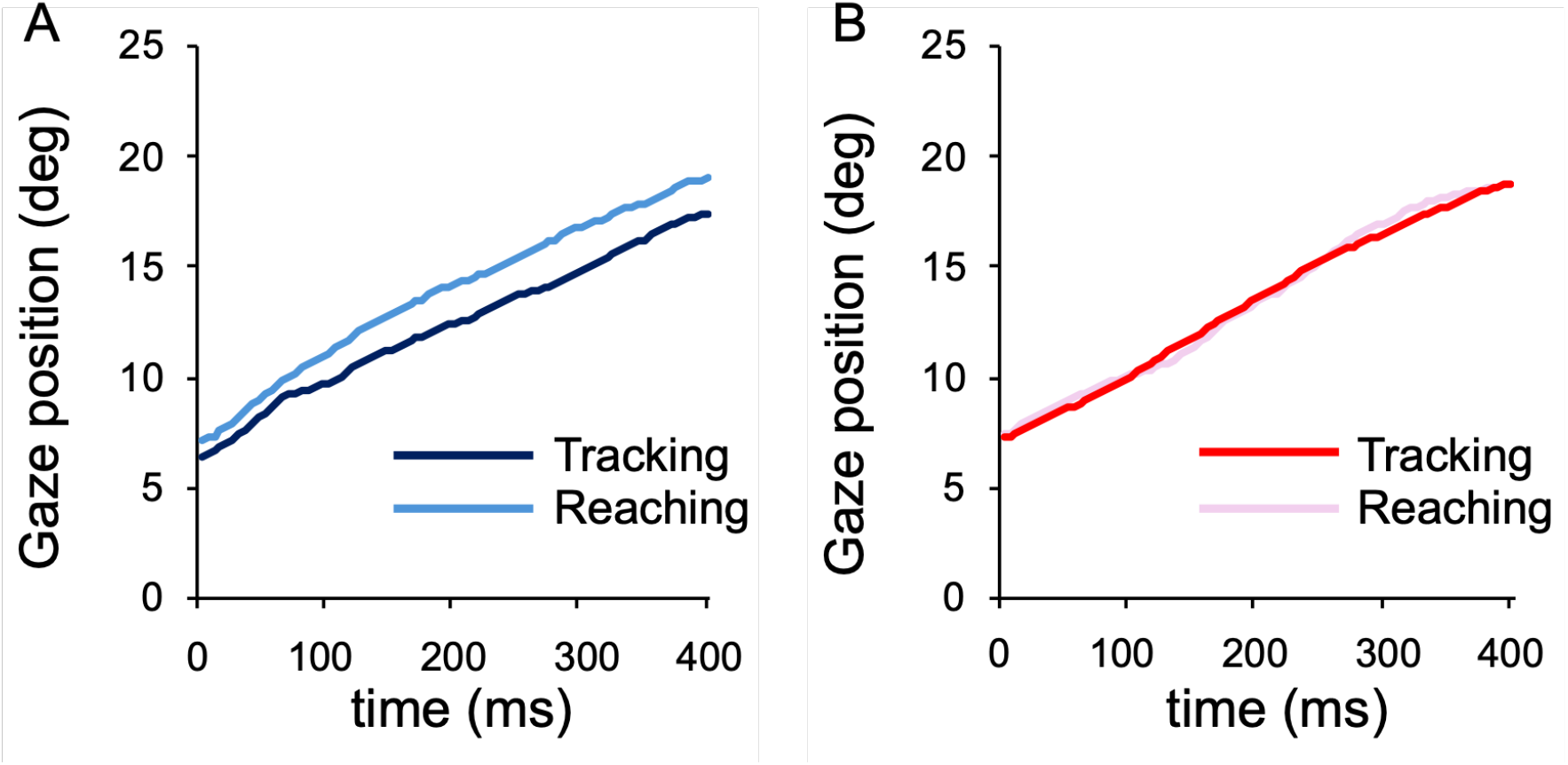
Representative examples of gaze position time series: (A) Trial-averaged gaze position of a representative novice participant. (B) Trial-averaged gaze position of a representative experienced participant. Each panel shows trial-averaged waveforms for the tracking task (dark blue or dark red) and reaching task (light blue or pink) over the 400 ms analysis window. t = 0 indicates the onset of the analysis window (20 ms after the offset of the initial saccade).

**Fig. 6.**
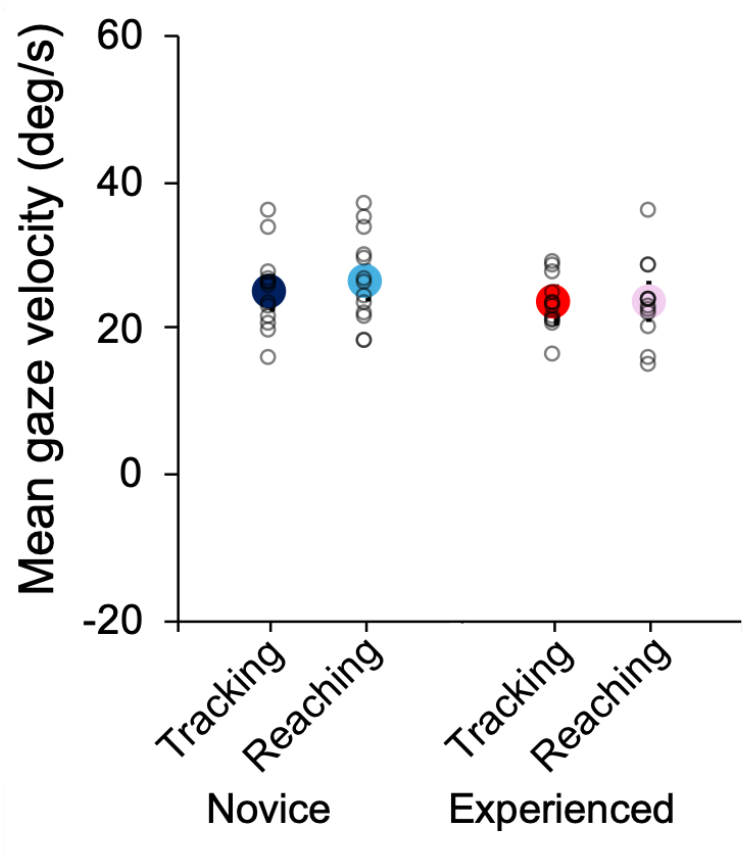
Comparison of mean gaze velocity across tasks and groups: Filled circles represent estimated marginal means; error bars indicate standard error. Open circles represent individual participant means, each averaged across that participant’s valid trials. Dark blue: novice group, tracking task; light blue: novice group, reaching task; dark red: experienced group, tracking task; pink: experienced group, reaching task.

### 3.3 Task- and experience-dependent changes in temporal gaze-head coordination

LMM results for the maximum cross-correlation coefficient between gaze and head velocity showed a significant main effect of task (*F* (1, 604.69) = 48.57, *p* < .001, η^2^p = .074). The cross-correlation coefficient was significantly larger in the reaching task (0.573 ± 0.025) than in the tracking task (0.494 ± 0.025) (Fig. 7A). The main effect of group was not significant (*F* (1, 23.16) = 0.31, *p* = .583, η^2^p = .013). The Task × Group interaction was significant (*F* (1, 604.69) = 5.05, *p* = .025, η^2^p = .008), indicating that experienced players exhibited a greater increase in the maximum cross-correlation coefficient from the tracking to the reaching task than the novice group. Simple effects tests for task confirmed that the cross-correlation coefficient was significantly larger in the reaching task than in the tracking task in both the novice group (tracking: 0.520 ± 0.034, reaching: 0.574 ± 0.034; *p* < .001) and the experienced group (tracking: 0.467 ± 0.036, reaching: 0.572 ± 0.036; *p* < .001). Simple effects tests for group showed no significant between-group differences in either the tracking task (*p* = .302) or the reaching task (*p* = .977).

LMM results for the lag time at the maximum cross-correlation between gaze and head velocity showed a significant main effect of group (*F* (1, 23.85) = 4.49, *p* = .045, η^2^p = .158). Because negative values indicate that gaze velocity preceded head velocity, the novice group showed a significantly longer gaze-leading interval (−24.19 ± 7.24 ms) than the experienced group (−1.54 ± 7.86 ms) (Fig. 7B). The main effect of task (*F* (1, 611.14) = 0.73, *p* = .394, η^2^p = .001) and the Task × Group interaction (*F* (1, 611.14) = 0.25, *p* = .615, η^2^p = .000) were not significant.

**Fig. 7.**
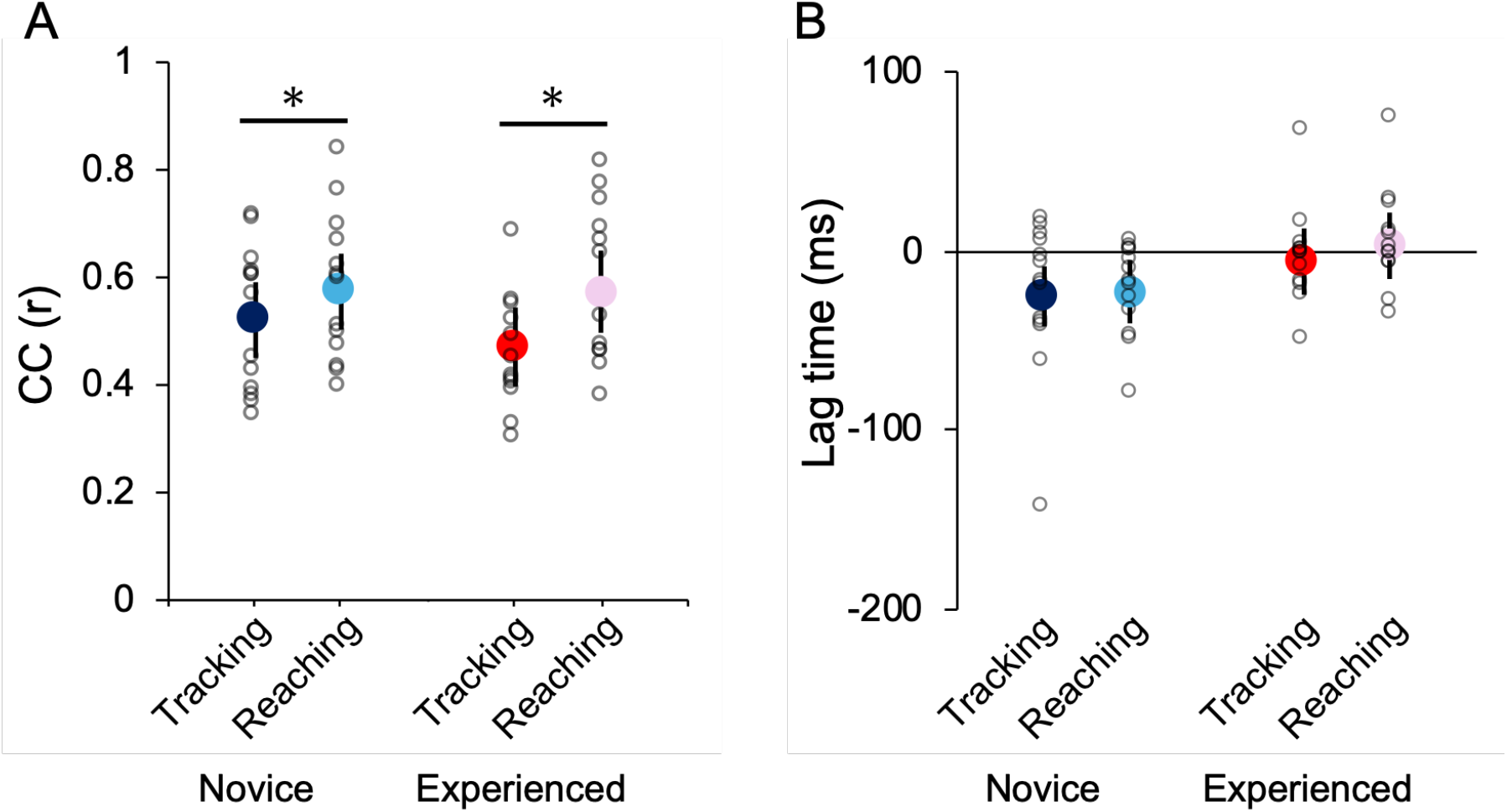
Cross-correlation between gaze and head velocity: (A) Comparison of cross-correlation coefficient (CC) between gaze and head velocity across tasks and groups. (B) Comparison of lag time between gaze and head velocity across tasks and groups. Negative values indicate that gaze leads head movement. Filled circles represent estimated marginal means; error bars indicate standard error. Open circles represent individual participant means, each averaged across that participant’s valid trials. Dark blue: novice group, tracking task; light blue: novice group, reaching task; dark red: experienced group, tracking task; pink: experienced group, reaching task. The horizontal line in panel B indicates zero lag. Negative lag time indicates that gaze velocity preceded head velocity.

## 4. Discussion

### 4.1 Constancy of gaze velocity

Gaze velocity did not differ significantly across tasks or groups. This result can be interpreted by decomposing gaze velocity into head velocity in space and eye velocity in the head. Despite opposite directional changes, greater eye velocity in the tracking task and greater head velocity in the reaching task offset each other, maintaining gaze velocity constant. Stable tracking of an approaching ball requires gaze movement to match ball trajectory, and the constancy of gaze velocity is therefore considered functionally important in meeting this demand. At the same time, the head-eye coordination pattern underlying this constancy differed as a function of task demands, and this is examined in detail in the following sections.

### 4.2 Changes in head and eye velocity as a function of task demands

By decomposing gaze velocity into head and eye components, the present study revealed task-dependent coordination changes that cannot be detected from gaze velocity alone. As shown in Section 4.1, gaze velocity did not differ significantly between tasks; however, decomposing gaze into head and eye components revealed that, compared with the tracking task, the reaching task led to significantly larger head velocity and significantly smaller eye velocity, reflecting opposite directional changes. These results indicate that task demands were associated with changes in the relative contributions of the head and eyes to gaze control, and that the coordinative mechanism underlying gaze velocity constancy cannot be detected through gaze-level analysis alone.

One possible explanation for the larger head velocity in the reaching task is that the head became more actively involved in predicting the ball’s landing position. Gaze control has been shown to contribute to predicting the ball landing position during catching tasks [22]; however, the present reaching task did not involve actual catching, and the analysis window was limited to the tracking phase. Nevertheless, the fact that head velocity was already elevated during the tracking phase suggests that the motor demands specific to the reaching task influenced head movement even before the reaching phase began.

Accurate hand movement to the ball’s landing position in the reaching task requires the transformation of visual information into body-centered coordinates. Eye and head orientation signals have been shown to be essential for integrating visual input with proprioceptive coordinates [15,16,23]. The greater head involvement observed in the reaching task may therefore have functioned to enhance the accuracy of this coordinate transformation.

Furthermore, reaching accuracy toward a spatially fixed target during body movement has been shown to be maintained through vestibular feedback [24]. In the present reaching task, the increase in head velocity may have augmented semicircular canal input, thereby contributing to the maintenance of spatial orientation accuracy.

In the tracking task, eye velocity was significantly larger than in the reaching task. This reflects the comparatively smaller head velocity in the tracking task and can be interpreted as the eyes compensating for the reduced head contribution to maintain gaze velocity. This pattern of opposite changes also represents a finding that cannot be detected by gaze analysis alone and further demonstrates the task-dependent coordination structure revealed by decomposing gaze into head and eye components.

### 4.3 Effects of ball sport experience on task-dependent modulation of eye velocity

Contrary to our hypothesis that novice players would show greater head and eye velocity than experienced players, no significant main effect of group was observed for either head velocity (*p* = .054) or eye velocity (*p* = .157). This suggests that ball sport experience does not substantially influence the magnitude of head and eye velocity itself. Although the Task × Group interaction for head velocity was not significant (*p* = .182), the Task × Group interaction for eye velocity was significant. This interaction indicates that the experienced group showed greater task-dependent change in eye velocity than the novice group. In other words, the experienced group more flexibly adjusts the relative contributions of eye movements in response to task demands. These results suggest that ball sport experience is associated with greater flexibility in head-eye coordination rather than consistently different movement patterns. Subsequent simple effects tests revealed that the simple effect of task was significant in both the novice and experienced groups, whereas the simple effect of group was not significant in either task. Previous studies have primarily compared gaze strategy within a single task and reported differences between experienced and novice players [17,18,25–28]. The present study extends these findings by showing that expertise is characterized not as differences in gaze strategy within a single task, but as the magnitude of eye velocity change across tasks with different behavioral demands.

### 4.4 Task- and experience-dependent changes in temporal gaze-head coordination

The maximum cross-correlation coefficient between gaze and head velocity was significantly larger in the reaching task than in the tracking task. Given the significant Task × Group interaction, simple effects tests were conducted, revealing that the simple effect of task was significant in both the novice and experienced groups, with cross-correlation coefficients being greater in the reaching task. The simple effect of group was not significant in either task.

The larger cross-correlation coefficient in the reaching task indicates that head-gaze coordination was stronger under conditions of greater task demand. This is consistent with the greater head involvement and can be interpreted as reflecting closer coordination between head and gaze for predicting the ball’s landing position. Furthermore, it has been shown that the combination of eye and head movements is determined by an optimal control principle that minimizes the impact of noise on motor performance [29], suggesting that the strengthened head-gaze coordination observed in the reaching task may reflect the selection of an optimal coordination pattern in response to increased task demands.

The significant Task × Group interaction suggests that the effect of task on gaze-head coupling differed between groups. Specifically, experienced players exhibited a greater increase in the maximum cross-correlation coefficient from the tracking to the reaching task than the novice group.

### 4.5 Experience-dependent differences in lag time

The main effect of group on lag time was significant: the novice group (−24.19 ms) exhibited a significantly longer gaze-leads-head interval than the experienced group (−1.54 ms). The near-zero lag observed in the experienced group might superficially resemble the timing characteristics of the VOR, in which compensatory eye movements occur opposite to head movements with minimal delay. However, if the near-zero lag were attributable to the VOR, eye movements would be expected to occur in the opposite direction to head movement, resulting in negative eye velocity values. Only a small number of participants in either group showed negative eye velocity (Fig. 4B), suggesting that VOR suppression was largely achieved in both groups. Thus, the group differences in lag time cannot be explained by VOR alone.

A possible interpretation is that the novice group exhibited a gaze-leading coordination pattern in which gaze precedes head movement, with the eyes first tracking the ball while the head followed with delay. In contrast, the near-zero lag observed in the experienced group suggests that eye and head movements show a synchronized coordination pattern, enabling the two effectors to work together in maintaining stable gaze on the approaching ball. Such coordinated timing is consistent with predictive, feedforward control, in which anticipatory motor commands are generated before visual feedback becomes available. Predictive smooth pursuit mechanisms driven by internal signals have been demonstrated even in the absence of visual feedback [30–33], indicating that eye movements can be generated in a feedforward manner without relying on continuous visual input. The present findings extend these observations by suggesting that experienced players not only engage predictive control of eye and head movements, but also temporally coordinate these movements to optimize ball tracking.

Previous studies examining head and eye movements during ball-catching tasks have primarily focused on describing the relative contributions of the eye and head to gaze shifts and the spatial patterns of their coordination [10,18,22]. More broadly, reviews of eye movements in manual interception tasks have similarly emphasized retinal and spatial contributions over temporal coordination patterns [34,35]. However, the temporal coordination between head and gaze velocity, as quantified through time-series analysis, has not been extensively addressed. The approach used in the present study offers a means of characterizing the temporal relationship between head and gaze movements and suggests the potential for a more precise description of experience-dependent differences in visuomotor coordination.

## 5. Limitations

Although the present study identified experience-dependent differences in head-eye coordination under controlled experimental conditions, future studies should examine whether these findings generalize to athletes from different ball sports and to more ecologically valid interception tasks involving actual catching. In addition, incorporating more detailed measures of expertise, such as competitive level and training history, may further clarify the relationship between motor experience and head-eye coordination.

## 6. Conclusion

By decomposing gaze into head and eye components during ball tracking, the present study showed that head-eye coordination is flexibly reorganized according to task demands and ball sport experience. Although gaze velocity remained unchanged across conditions, head and eye velocities changed in opposite directions between tasks, indicating that gaze constancy is maintained through coordinated adjustments of the two components. Experienced players exhibited greater task-dependent modulation of both eye velocity and gaze-head cross-correlation, and also showed a consistently smaller gaze-head lag than novice players regardless of task. These results suggest that ball sport experience shapes two distinct aspects of head-eye coordination: task-dependent flexibility and predictive gaze-head control. Taken together, our findings provide insight into the characterization of visuomotor coordination that supports skilled performance in open-skill sports.

## CRediT authorship contribution statement

**Saya Ogino**: Conceptualization, Methodology, Software, Formal analysis, Investigation, Data curation, Writing – original draft, Writing – review & editing, Visualization. **Tomohiro Kizuka**: Conceptualization, Funding acquisition, Methodology, Supervision. **Seiji Ono**: Conceptualization, Funding acquisition, Methodology, Writing – review & editing, Supervision

## Data availability

Data will be made available on request.

